# Body size and climate as predictors of plumage colouration and sexual dichromatism in parrots

**DOI:** 10.1101/2020.05.21.107920

**Authors:** Luisana Carballo, Kaspar Delhey, Mihai Valcu, Bart Kempenaers

## Abstract

Psittaciformes (parrots, cockatoos and lorikeets) comprise one of the most colourful clades of birds. Their unique pigments and cavity nesting habits are two potential explanations for their colourful character. However, plumage colour varies substantially between parrot species and sometimes also between males and females of the same species. Here, we use comparative analyses to evaluate what factors correlate with colour elaboration, colour diversity and sexual dichromatism. Specifically, we test the association between different aspects of parrot colouration and (1) the intensity of sexual selection and social interactions, (2) variation along the slow-fast life-history continuum and (3) climatic variation. We show that larger species and species that live in warm environments display more elaborated colours, yet smaller species have higher levels of sexual dichromatism. Larger parrots tend to have darker and more blue and red colours. Parrots that live in humid environments are darker and redder, whilst species inhabiting warm regions have more blue plumage colours. In general, the variables we considered explain small to moderate amounts of variation in parrot colouration (up to 20%). Our data suggest that sexual selection may be acting more strongly on males in small, short-lived parrots leading to sexual dichromatism. More elaborate colouration in both males and females of the larger, long-lived species with slow tropical life-histories suggests that mutual mate choice and reduced selection for crypsis may be important in these species, as has been shown for passerines.

## Introduction

Birds show great diversity in plumage colour and many studies have aimed to explain the proximate and ultimate mechanisms behind this diversity (Baker & Parker, 1979; Dale, Dey, Delhey, Kempenaers, & Valcu, 2015; Delhey, 2017, 2018; Hill & McGraw, 2006; Miller, Leighton, Freeman, Lees, & Ligon, 2019; Taysom, Stuart-Fox, & Cardoso, 2011). Among birds, Psittaciformes – parrots, cockatoos and lorikeets (from now on collectively called parrots) – show some of the most striking plumage colouration (Berg & Bennett, 2010; Delhey, 2015). However, the evolutionary forces underlying their colourful character remain poorly understood (Berg & Bennett, 2010). It has been argued that parrots are colourful because they can synthesise and deposit red and yellow psittacofulvin pigments in their feathers, which are unique to parrots (McGraw & Nogare, 2004; Stradi, Pini, & Celentano, 2001). Because these pigments are synthesised endogenously, parrots might be able to deposit higher concentrations and display more intense colours compared with other bird species that can only obtain carotenoids (to produce yellow to red colours) through their diet (Delhey, 2015). Psittacofulvins, in combination with melanin pigments and feather microstructural components (which produce structural colours such as blue), enable parrots to display colours that encompass a large proportion of the entire avian colour gamut (Berg & Bennett, 2010; Delhey, 2015). In addition, most parrots breed in cavities, which are safe nesting sites that provide protection to parents and offspring from predators (Martin & Pingjun Li, 1992), potentially removing the need to be cryptic at the nest. Parrots, both males and females, are indeed more colourful than expected for their species richness (Delhey, 2015) and many species are mutually ornamented (Berg & Bennett, 2010).

Parrots are generally colourful, but also show great colour variation among species. For example, some cockatoo species are monochromatic and entirely white, whilst the Eclectus parrot (*Eclectus roratus*) is highly sexually dichromatic, with males being mainly green and females bright red and blue. The selective forces behind this substantial variation in colour elaboration and sexual dichromatism within parrots (Delhey, 2015; Delhey & Peters, 2017; Taysom *et al*., 2011) are not yet well understood (Berg & Bennett, 2010).

Ornamental traits might be used in competitive interactions or in sexual displays. For this reason, many studies have explored how sexual and social interactions may have driven plumage colour evolution (Dale *et al*., 2015; Dunn, Whittingham, & Pitcher, 2001; Miller *et al*., 2019; Møller & Birkhead, 1994; Owens & Hartley, 1998; Rubenstein & Lovette, 2009). Colour traits can be favoured by sexual selection if the expression of the trait increases the reproductive success of individuals by gaining more access to mates, or by social selection if their expression is critical in the competition for social status or access to resources such as food or territories (West-Eberhard, 1983).

The intensity of sexual selection, as found in polygynous species, correlates with the occurrence of multiple ornaments (Møller & Pomiankowski, 1993) and sexual dichromatism in birds (Dale *et al*., 2015; Dunn *et al*., 2001). In lizards, two proxies for sexual selection intensity (sexual dimorphism in size and colour) correlate positively with colour diversity, i.e. the different colours and patterns an individual displays (Chen, Stuart-Fox, Hugall, & Symonds, 2012). Additionally, bird species with high levels of extra-pair paternity presumably experience stronger sexual selection and also show higher levels of sexual dichromatism (Møller & Birkhead, 1994; Owens & Hartley, 1998). A large-scale comparative analysis in passerines showed that sexual selection is the strongest predictor of sexual dichromatism (Dale *et al*., 2015).

Colour ornamentation may have also evolved in response to the selective pressures of complex social interactions (Heinsohn, Legge, & Endler, 2005; Santana, Alfaro, Noonan, & Alfaro, 2013). For group living species, such as parrots, it might be advantageous to effectively signal status, age or identity (Bridge, Hylton, Eaton, Gamble, & Schoech, 2008; Dale *et al*., 2015), which may be easier to achieve with multiple signals (e.g. with higher colour diversity). Support for this idea comes from primates, where the complexity of facial markings is correlated with gregariousness (Santana *et al*., 2013). Further support comes from a study on the Eclectus parrot, showing that the extreme scarcity of suitable nest cavities (~1 per square kilometre) has intensified intrasexual competition (Heinsohn *et al*., 2005). Females spent most of their time protecting their nest (for around 11 months a year) and they may kill each other in disputes over tree hollows (Heinsohn *et al*., 2005). Thus, Heinsohn *et al*. (2005) suggested that the expression of conspicuous colours in females is a consequence of the need to display cavity ownership.

With a few exceptions, the mating system of parrots is social monogamy (Toft & Wright, 2015), which implies lower levels of sexual selection. However, a recent study showed considerable variation in sperm length in parrots, with sexually dichromatic and gregarious species having longer sperm (Carballo *et al*., 2019). This suggests that some parrots experience higher levels of sperm competition, for example due to increased opportunities for extra-pair mating when pairs nest in close proximity (Møller & Birkhead, 1993). We can thus ask whether variation in sexual dichromatism, colour elaboration and colour diversity are linked to indicators of the intensity of sexual selection in parrots.

The intensity of sexual selection may also depend on the species’ life-history strategy (Winemiller, 1992). Given that the lifespan of parrots ranges from 8.5 to 100 years (Wasser & Sherman, 2010), one can explore whether the slow-fast life-history continuum is linked to parrot plumage colouration. In general, parrots form long-lasting pair bonds and the formation of such bonds may take time (Toft & Wright, 2015). Smaller parrot species experience a higher turnover of mates (Toft & Wright, 2015), which might be related to the higher mortality rate associated with smaller body size (de Magalhaes, Costa, & Church, 2007; Wasser & Sherman, 2010). Consequently, the expression of sexually selected traits that help speed up the selection of mates could be more beneficial for females in species with lower adult survival if it reduces the time needed to identify a suitable male and form a pair bond. On the other hand, long-lived species with long-lasting pair bonds might experience mutual mate choice, linked to higher parental investment in both sexes (Kokko & Johnstone, 2002). In such cases, both males and females are expected to be more elaborately coloured. Larger species also experience reduced predation risk, a factor that may explain why males and females of larger passerine species have more elaborated colours (Dale *et al*., 2015). Furthermore, the slow-fast life-history continuum is related to extra-pair paternity: species with higher adult mortality rates and larger clutch sizes have higher levels of extra-pair paternity (Arnold & Owens, 2002).

Different studies have evaluated how abiotic factors affect bird plumage colour evolution and a variety of hypotheses have been proposed to explain colour variation both within and across avian taxa (Dale *et al*., 2015; Miller *et al*., 2019; Ribot, Berg, Schubert, Endler, & Bennett, 2019). Previous studies showed that achromatic (light-to-dark) variation in birds is related to climate variables such as temperature and precipitation (Delhey, 2017, 2018, 2019; Heidrich *et al*., 2018; Miller *et al*., 2019; Pinkert, Brandl, & Zeuss, 2017; Ribot *et al*., 2019). Specifically, a negative relationship between melanin pigmentation and temperature has been reported in several taxa (Delhey, 2018; Heidrich *et al*., 2018; Pinkert *et al*., 2017), in support of the thermal melanism hypothesis (Clusella Trullas, van Wyk, & Spotila, 2007). This eco-geographical rule proposes that darker animals inhabit colder environments, presumably for thermoregulation reasons (Clusella Trullas *et al*., 2007; Delhey, 2018). Similarly, Gloger’s rule suggests a positive association between melanin pigmentation and precipitation (Delhey, 2017, 2019; Gloger, 1833), but the adaptive function of the link between darker colours and precipitation is not yet clear (Burtt & Ichida, 2004a; Delhey, 2017; Zink & Remsen, 1986).

In summary, different factors may affect plumage colouration and sexual dichromatism. Therefore, to better understand what factors might explain interspecific variation in colour elaboration, colour diversity and sexual dichromatism, it is important to consider multiple variables simultaneously. So far, few studies on plumage colouration have considered multiple variables. Dale *et al*. (2015) used comparative analyses to explore the effects of multiple traits on plumage colour in passerines. Specifically, this study suggests that the evolution of plumage colour and sexual dichromatism are mainly driven by sexual selection and life-history traits, with stronger effects on female than on male colour. Both males and females are more colourful in larger species and in species with tropical life histories (i.e. small clutch size, low seasonality habitats), whilst sexual dichromatism was higher in smaller species and in species with male-biased sexual selection.

Here, we ask what factors affect plumage colouration in parrots.We quantified achromatic and chromatic colour variation among all 398 species of the order Psittaciformes based on colour plates, and computed estimates of colour elaboration, colour diversity and sexual dichromatism. Our study had three main aims. (1) To test whether indicators of the intensity of sexual selection and social interactions relate to variation in plumage colouration in parrots. We predict higher sexual dichromatism, and higher colour elaboration and colour diversity in males in species that (a) show stronger male-biased sexual size dimorphism and (b) breed at higher densities (i.e. are gregarious). (2) To test whether the slow-fast life-history continuum is associated with plumage colour variation in parrots. We predict higher sexual dichromatism, and higher colour elaboration and colour diversity in males in species that (a) have smaller body size (because body size correlates positively with longevity; Wasser & Sherman, 2010) and (b) lay larger clutches. We predict lower sexual dichromatism but higher colour elaboration and colour diversity in both males and females (mutual ornamentation) in species that (c) have large body size and (d) lay smaller clutches. (3) To test whether parrots follow Gloger’s rule and the thermal melanism hypothesis. If so, we predict that (a) darker species inhabit more humid and colder environments and (b) darker species inhabit densely forested rather than open habitat types (because the former are typically more humid).

## Material and methods

### Plumage colour scores

We compiled digital images of colour plates of both sexes for each of the 398 extant parrot species illustrated in the *Handbook of the Birds of the World Alive* (HBW Alive, del Hoyo *et al*., 2017). We imported the images into *Adobe Photoshop* (Adobe Inc. San Jose, CA), cropped them to remove the background colour and all bare parts of the birds, thus keeping only the body regions covered by plumage, and saved them as PNG files. Subsequently, we delineated 12 body patches (nape, crown, forehead, throat, upper breast, lower breast, shoulder, secondary coverts, primary coverts, secondaries, primaries and tail) for each sex and extracted RGB (red, green, blue) colour values from 400 randomly chosen pixels in each patch using the R package “colorZapper” v.1.4.4 (Valcu & Dale, 2014). Even though the different body patches differed in size, we randomly selected 400 pixels from each patch, because body regions may vary in signalling importance. For the monochromatic species (i.e. when one plate is shown to represent both male and female), the colour values were randomly extracted twice (once for the male and once for the female). In some cases, the plates of one of the sexes did not show the entire body, hence the colour values of the missing body patches were extracted from the plate of the other sex. When multiple subspecies were illustrated, the nominate species was scored. Finally, we calculated mean R, G and B values for each patch, sex and species. We transformed these mean values to CIELAB coordinates (Tkalčič & Tasič, 2003) using the R package “colorspace” v.1.4-1 (Zeileis *et al*., 2019). There are three CIELAB coordinates: (1) *L*, colour lightness, represents the achromatic channel (black = 0, white = 100, Figure 1a), (2) *a*, the chromatic channel between green (low values) and red (high values) (Figure 1b) and (3) *b*, the chromatic channel between blue (low values) and yellow (high values) (Figure 1c). We used the CIELAB coordinates to compute the following colour variables:

a. *Colour elaboration score*, obtained by computing the Euclidean distance between each plumage patch and the centroid of the entire sample (joint average for *L, a*, and *b*). These values were averaged in each species, separately for males and females. Highly elaborate colours (in this case, red, blue and yellow) are those that differ more from the average colour (here: greenish brown) (Figure 1d). This index of colour elaboration yields a similar classification of elaborate colours as the one used in Dale *et al*. (2015) (compare Figure 1d with Figure S2 in Dale *et al*., 2015).
b. Sexual differences in colouration, computed in two ways: (i) *Sexual dichromatism*, as the Euclidean distance in CIELAB space between homologous patches in males and females averaged across all patches for each species (Figure 2a), and (ii) *sexual difference in colour elaboration*, as the average difference in colour elaboration between males and females (Figure 2b). The first index (i) estimates the absolute difference in colouration between males and females irrespective of whether males or females are more ornamented. The second one (ii) indicates whether it is males or females that have more elaborated colours. Note that if males and females have different colours but with the same level of elaboration (e.g. red and blue) this index will score low.
c. Three overall plumage colour scores for each sex and species by calculating average values for *L, a*, and *b* of all 12 body patches (Figure 1a-c, and see Figure S1 for more details of the raw colour distribution of each body patch). This allows us to assess whether explanatory variables favour the evolution of certain types of colours over others (e.g. red over green, light over dark). The downside of this approach is that species that harbour a wide range of colours may end up with intermediate average values of *L*, *a* or *b*.
d. Finally, we estimated *colour diversity*, computed as the Euclidean distance between each plumage patch and the species-specific (rather than that of the entire sample as in (a)) centroid (joint average for *L*, *a*, and *b* of all plumage patches of each species). This measure indicates whether a species has many different colours or is rather uniformly coloured.

**Figure 1.**
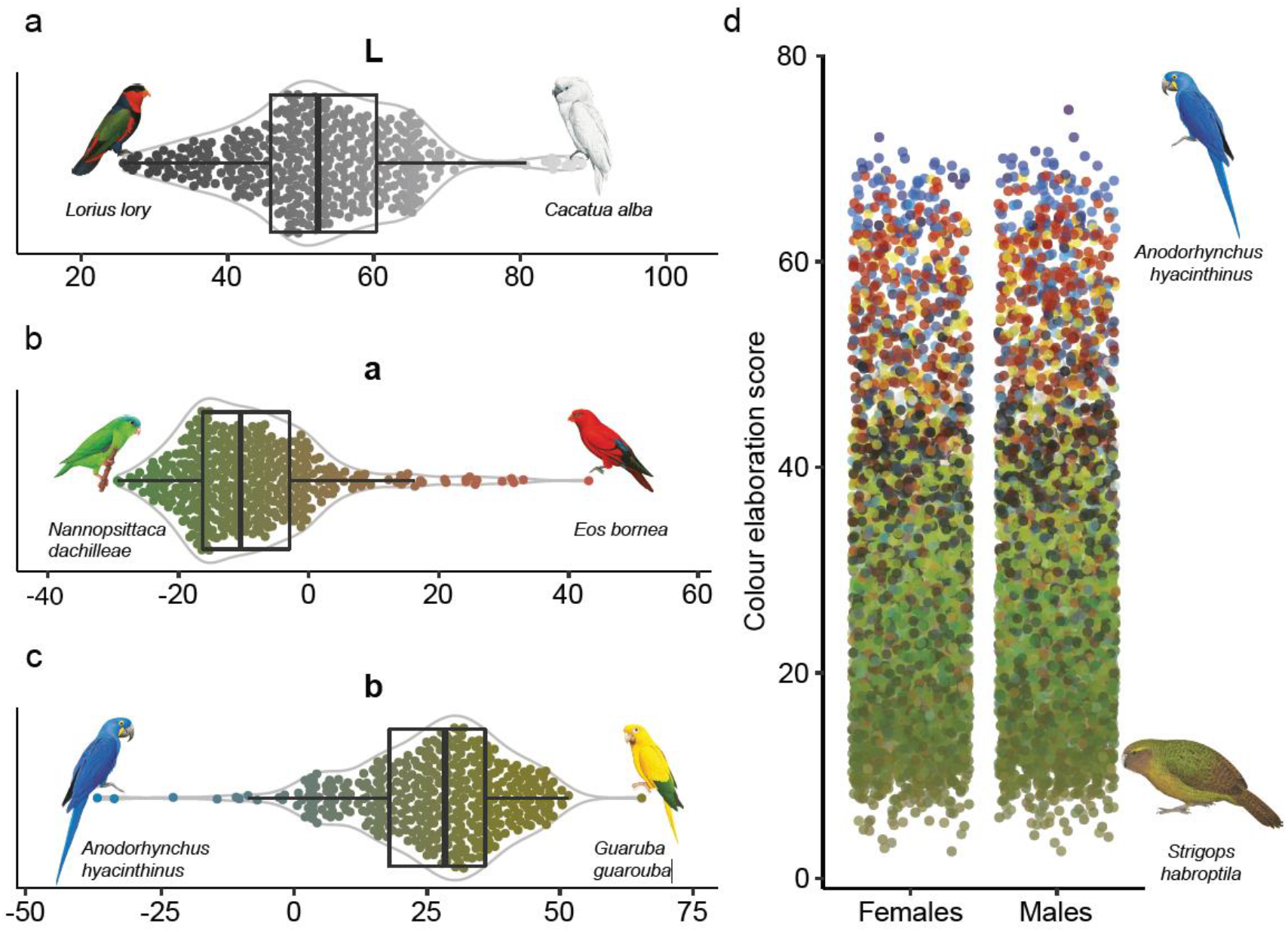
Illustration of the plumage colour scores for 398 parrot species. **a** *L*-score distribution showing dark to light colours, **b** *a*-score distribution showing green to red colours, **c** *b*-score distribution showing blue to yellow colours, and **d** colour elaboration score of females and males showing the distribution from the average colour (greenish brown) to highly elaborate colours such as red, blue, yellow, black and white. Illustrations in each panel represent the species that have the minimum and maximum scores for each variable. **a**-**c**, shown are box plots with median (vertical line) and interquartile range (box), and violin plots (grey lines) showing the probability density of the data. The dots in **a-c** represent the colour of each species for each colour coordinate (averaged across 12 body patches). To show the colour score of each species on the *L*, *a* and *b* coordinates separately, variation in the focal colour coordinate is shown while the other two colour coordinates were fixed (**a**, *a* = 0, *b* = 0; **b***, L* = 50, b = 26.4 (mean score for all species); **c**, *L* = 50, *a* = −8.8 (mean score for all species)). Illustrations © Lynx Edicions.

**Figure 2.**
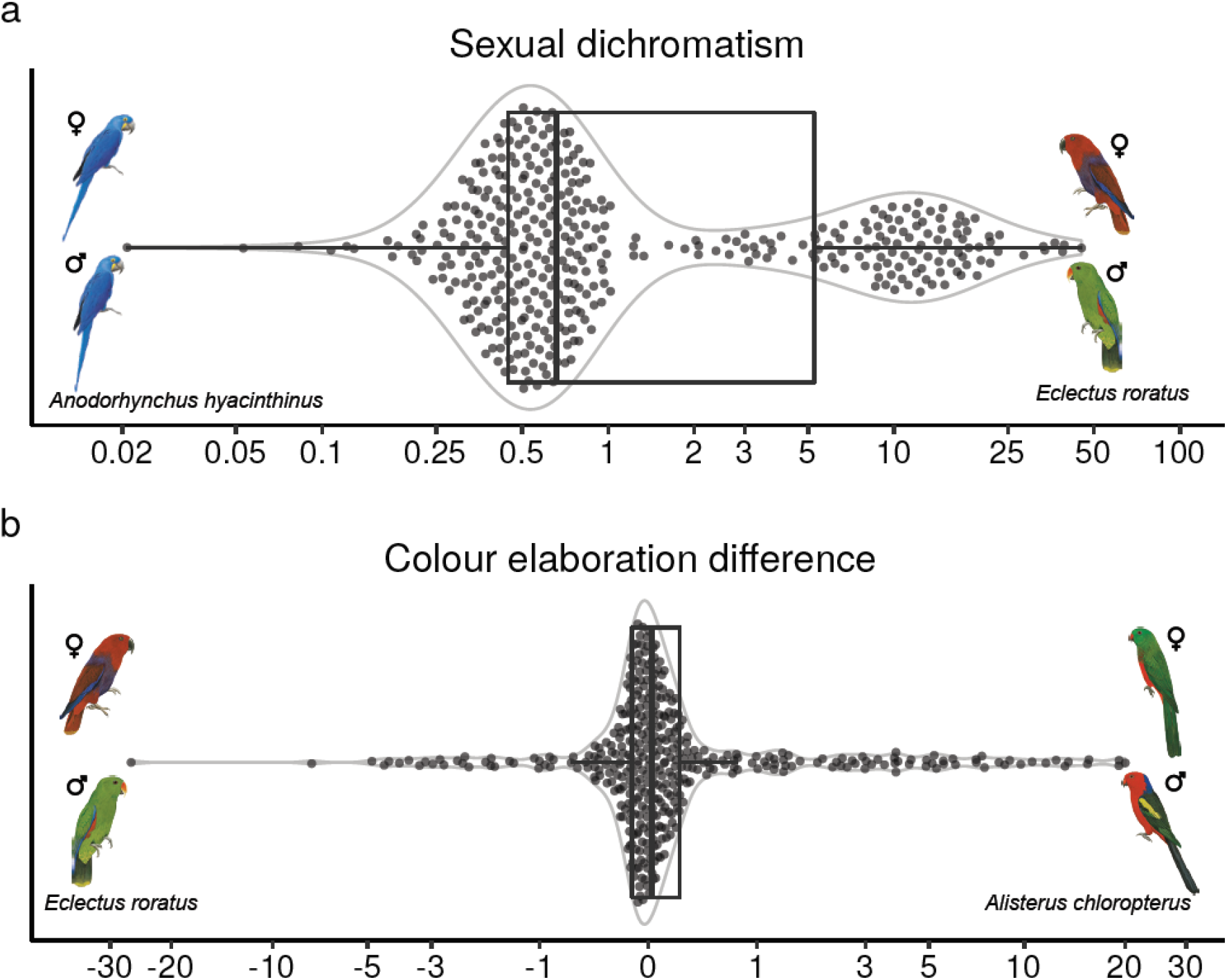
Illustration of sexual differences in colouration for 398 parrot species. **a** Distribution of the sexual dichromatism score, **b** distribution of sexual differences in colour elaboration. X-axes scales are log10 transformed and log10-modulus transformed (sign(x)*log10(abs(x)+1), John and Draper, 1980) for negative values. Illustrations in each panel represent the species that have the minimum and maximum scores for each variable. Shown are box plots with median (vertical line) and interquartile range (box), and violin plots (grey lines) showing the probability density of the data. Illustrations © Lynx Edicions.

The colour plates in the HBW have been painted to resemble real plumage colours as perceived by humans. To determine whether our estimates approximated those obtained using direct measurements of plumage, we used reflectance measurements obtained from 51 species of Australian parrots and cockatoos (Delhey, 2015; see Supplementary Information).

In general, all variables obtained from bookplates were positively correlated with estimates from reflectance spectra (all p < 0.001). Colour elaboration scores showed the weakest correlations (males: r = 0.53, females: r = 0.67), followed by differences in colour elaboration between males and females (r = 0.60), colour diversity (males: r= 0.83, females: r = 0.74) and sexual dichromatism (r = 0.86). *L* scores (which depict light-to-dark variation) were also positively correlated (males: r = 0.88, females: r = 0.89). It is harder to determine whether both chromatic coordinates in the CIELAB space, (*a* and *b*) correlate with the chromatic coordinates obtained from visual models (xyz, see Supplementary Information) because the latter do not necessarily align with the former. However, if both types of chromatic coordinates represent similar colours then we would expect that a linear combination of visual model chromatic coordinates (xyz) should predict chromatic coordinates (*a*, *b*) from bookplates. This was the case: xyz predicted substantial variation in *a* (males, R^2^ = 0.78; effects(SE): x = −0.277(0.521), y = - 2.406(0.246), z = 2.484(0.351); females, R^2^ = 0.85, x = −0.628(0.507), y = −3.123(0.258), z = 3.044(0.333) and *b* (males, R^2^ = 0.68, effects(SE): x = 2.931(0.831), y = 3.534(0.392), z = 1.114(0.558); females, R^2^ = 0.74, x = 3.257(0.898), y = 4.654(0.457), z = 0.031(0.588)). Thus, results obtained from bookplates should provide a reasonable approximation to colour variation measured on the plumage, as shown in other studies (Bergeron & Fuller, 2018; Dale *et al*., 2015).

### Measures of sexual selection and gregariousness

As a measure of the intensity of sexual selection, we calculated sexual size dimorphism (SSD) as *PC1_male body size_* – *PC1_female body size_* (see below). We scored gregariousness as a categorical variable (“yes” or “no”) according to information from the “breeding” section of the HBW Alive (del Hoyo *et al*., 2017). A species was classified as gregarious if the description suggested that the breeding pairs nest close together or if the species is described as colonial.

### Life-history traits

For each species, we estimated body size of males and females as the first principal component (PC1) from a PCA that included three body measurements: wing, tarsus and tail length. PC1 explained 65% of the variation in the data. We measured these traits for an average of 3.3 (*range*: 1-22) females and 3.6 (*range*: 1-23) males per species (N_species_ = 214) from individuals held at the Loro Parque Fundación (LPF), Tenerife, Spain. Species body size was estimated by calculating the average of male and female body size. For the species that were not present in the LPF collection, we compiled body measurements from the book *Parrots of the World* (Forshaw, 1978).

We obtained clutch size for each species from the HBW Alive (del Hoyo *et al*., 2017). As some species did not have clutch size data, we completed the database using LPF records from the 2012-2015 breeding seasons, by calculating the mean clutch size from 1-105 clutches per species (*mean* = 10.5), and using data available in the book *Parrots of the World* (Forshaw, 1978), and in the websites www.parrots.org and www.avianweb.com. The source of the body measurements and clutch size data for each species is given in the online repository.

### Environmental variables

We considered three environmental variables: habitat type, mean annual temperature (°C) and mean annual precipitation (mm). We scored habitat type as a categorical variable (1 = “open”, 2 = “mixed”, 3 = “forested”) using the description in the “habitat” section of the HBW Alive (del Hoyo *et al*., 2017). Following McNaught & Owens (2002), we classified habitat type as “open” for species that occur in habitats such as savannah, grassland, shrubland, forest edges, arid and eucalypt woodland or cliffs, as “forested” for species that occur in habitats such as forest, riverine forest, riparian forest, pine woodland, mangrove, evergreen lowland or wooded country, and as “mixed” for species that inhabit both “open” and “forested” habitat.

To estimate species-specific mean annual temperature and mean annual precipitation, we first obtained the extant breeding ranges for each parrot species using the database from BirdLife International’s species distribution maps (BirdLife International, 2018). We only considered the natural distribution of each species and hence removed any breeding ranges where they were introduced. We extracted the mean annual temperature and mean annual precipitation corresponding to the breeding ranges of each species using the high-spatial resolution CHELSA climate data (Karger *et al*., 2017a, 2017b). Breeding ranges and environmental rasters were re-projected to an equal-area (Mollweide) projection. Spatial analyses were performed with the R package “rangeMapper” v.0.3-7 (Valcu, Dale, & Kempenaers, 2012).

### Phylogeny

We extracted a sample of 1000 phylogenetic trees (the “Hackett” backbone, Hackett *et al*., 2008) for 351 parrot species from phylogenetic tree distributions available on *birdtree.org* (Jetz, Thomas, Joy, Hartmann, & Mooers, 2012; Jetz *et al*., 2014). We added the 47 Psittaciformes species missing in these phylogenies using the function *add.species.to.genus* in the R package “phytools” v.0.6-99 (Revell, 2012). This function finds the branch of the phylogenetic tree common to the corresponding genus and adds the missing taxon at a random position within this branch. A consensus tree was constructed with minimum clade frequency threshold of 0.5 (Rubolini, Liker, Garamszegi, Møller, & Saino, 2015) using the function *SumTrees* from the package “DendroPy” v.4.4.0 (Sukumaran & Holder, 2010).

### Statistical analysis

All statistical and spatial analyses were performed in R 3.6.2 (R Development Core Team, 2019). The variables sexual dichromatism and sexual difference in colour elaboration were log10 transformed and log10-modulus transformed (sign(x)*log10(abs(x)+1), John & Draper, 1980), respectively, for analyses. All variables were standardised by centring and dividing by one standard deviation.

To explore the effect of abiotic and biotic factors on plumage colour elaboration, sexual dichromatism and colour diversity across parrots, we used species-level phylogenetic linear models. These models were fitted with the R package “phylolm” v.2.6 (Ho & Ané, 2014) using the Pagel’s λ model (Pagel, 1999), which measures the strength of the phylogenetic signal. We ran separate models for our seven response variables, i.e. colour elaboration, sexual dichromatism, sexual difference in colour elaboration, colour diversity and the three plumage colour scores (*L*, *a* and *b*), and we considered body size (N = 357), clutch size (N = 290), habitat type (N = 398), mean annual temperature (N = 398), mean annual precipitation (N = 398), sexual size dimorphism (N = 357) and gregariousness (N = 350) as predictors in our analyses. First, we ran univariate models to explore the effect of each predictor separately, and allowing the use of the full dataset. For the 273 species for which all the predictors were available, we then ran a multiple predictor model to explore the effect of each predictor, whilst controlling for the others.

We estimated the proportion of variance explained by the phylogenetic linear models following Ives (2019) by using the function *R2.resid* in the R package “rr2” v.1.0.2 (Ives & Li, 2018). We calculated two R^2^ coefficients: (1) *R^2^_full_*: the total variance explained by the full model (both by phylogeny and fixed effects), and (2) *R^2^_fixef_*: the variance explained by the fixed effects only.

We ran species-level phylogenetic linear models for each of the 1000 phylogenies and we averaged the model coefficients. Additionally, we computed an inference interval as the 2.5 ^th^ - 97.5 ^th^ percentiles for p-values, Pagel’s λ and the two R^2^ coefficients. Therefore, the Pagel’s λ and the R^2^ coefficients inference intervals contain both the error of the distribution underlining the phylogenetic trees and the uncertainty of the taxonomy-based data imputation.

## Results

### Effects on plumage colouration

Both males and females of larger species and of species with smaller clutch size had more elaborated plumage colours. These effects were statistically significant in the single and multiple predictor models for body size (Figure 3, Table S1-S4), but the clutch size effect was statistically significant only in the single predictor models (Figure 3a). The lower effects and loss of significance of clutch size in the multiple predictor model (Figure 3b) might be due the intercorrelation between clutch size and body size (Figure S2). We also found that annual mean temperature had a positive effect on colour elaboration in both males and females; this effect was significant in the single and multiple predictor models (Figure 3, Table S1-S4).

**Figure 3.**
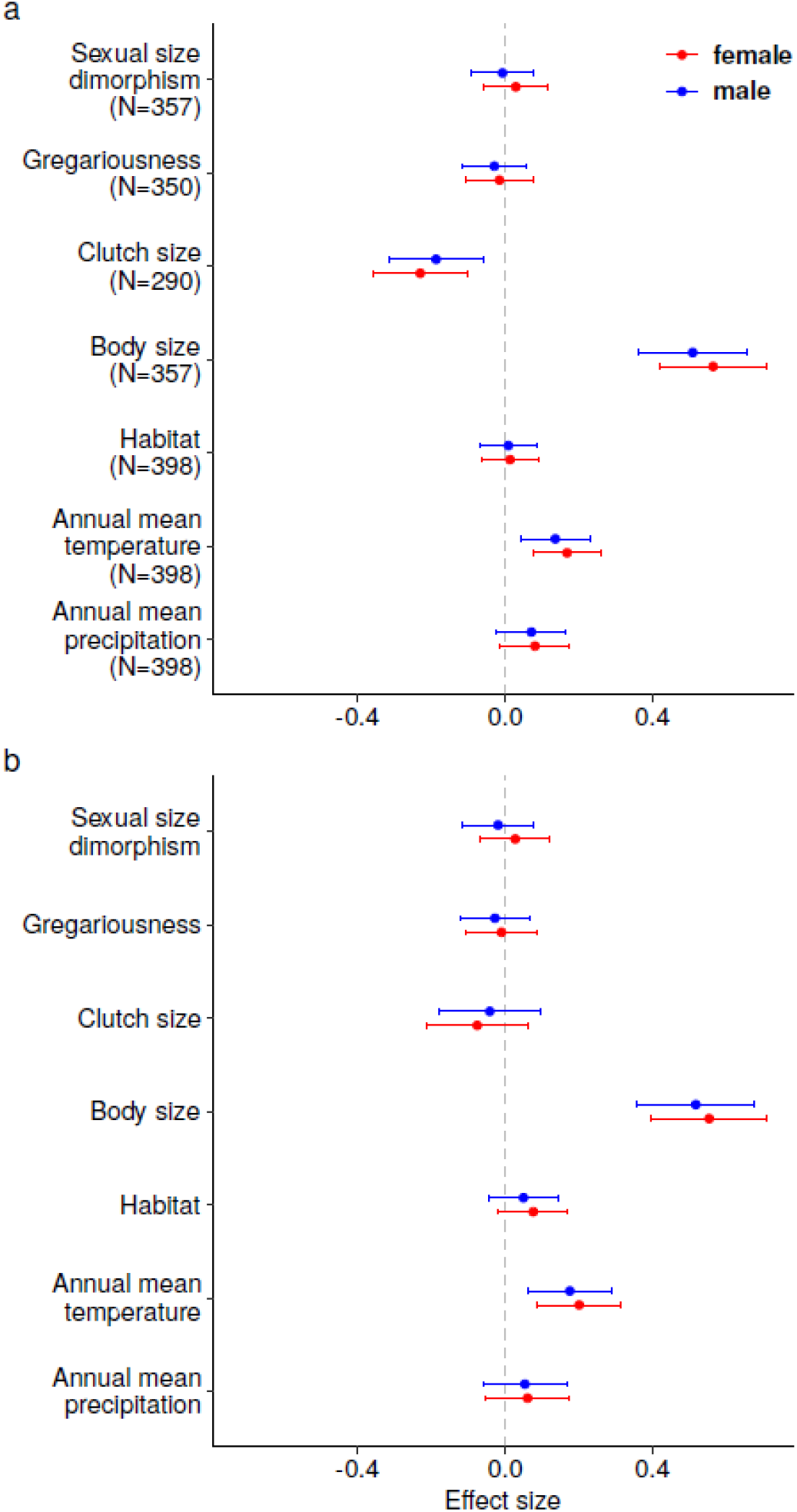
Effect sizes of predictors of colour elaboration based on **a** single predictor models and **b** a multiple predictor model (N = 273 species). Red denotes females and blue refers to males. Shown are the means of the model coefficients for the 1000 phylogenetic linear models and the corresponding 95% confidence intervals. N indicates the number of species included in the analyses (determined by data availability).

In both sexes, body size was significantly negatively associated with *L* and *b* scores and positively associated with *a* scores, both in the single predictor models (Figure 4a, Table S5 and S6) and in the multiple predictor model (Figure 4b, Table S7 and S8). These results suggest that males and females of larger species are darker, redder and more blue-coloured.

**Figure 4.**
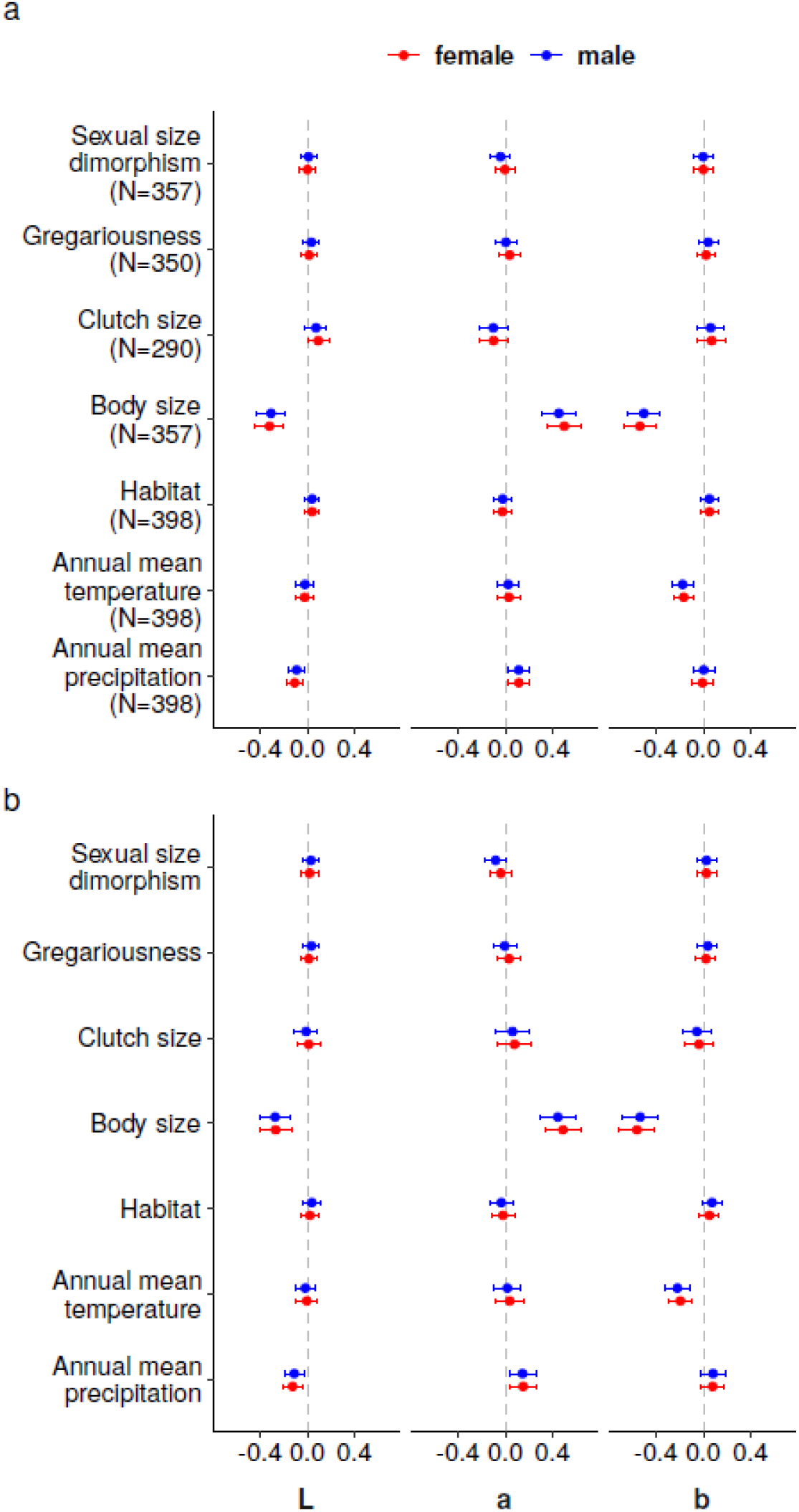
Effect sizes for each of the predictor variables on the three CIELAB colour coordinates (*L* = dark-to-light variation, *a* = green-to-red variation, *b* = blue-to-yellow variation), based on **a** single predictor models and **b** multiple predictor models (N = 273 species). Red denotes females and blue refers to males. Shown are the means of the model coefficients for the 1000 phylogenetic linear models and the corresponding 95% confidence intervals. N indicates the number of species included in the analyses (determined by data availability).

In both sexes, precipitation had a negative effect on *L* scores and a positive effect on *a* scores, whilst temperature had a negative effect on *b* scores in the single (Figure 4a, Table S5 and S6) and multiple predictor models (Figure 4b, Table S7 and S8). These results indicate that species that are darker and redder inhabit areas of higher mean annual precipitation, and that more blue-coloured species inhabit areas of higher mean annual temperature.

Habitat type did not have an effect on plumage colour in parrots (Figure 3 and 4, Table S1-S8), at least based on the data and classification used in this study.

### Effects on colour diversity

None of the predictors used in this study had a statistically significant effect on colour diversity in parrots, either in the single or in the multiple predictor models (Figure 5, Table S9-S12).

**Figure 5.**
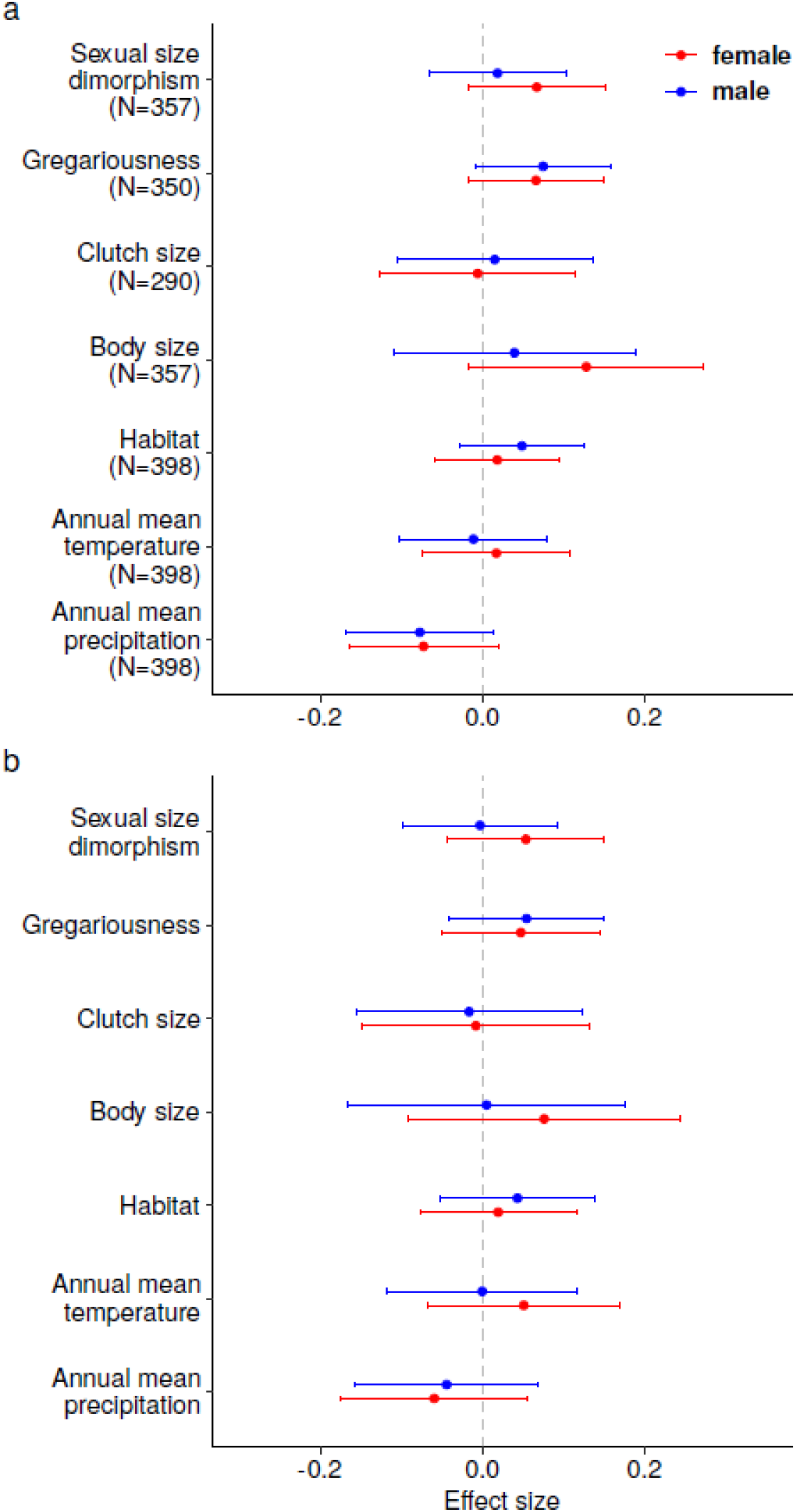
Effect sizes of predictors of colour diversity based on **a** single predictor models and **b** a multiple predictor model (N = 273 species). Red denotes females and blue refers to males. Shown are the means of the model coefficients for the 1000 phylogenetic linear models and the corresponding 95% confidence intervals. N indicates the number of species included in the analyses (determined by data availability).

### Effects on sexual differences in colouration

The single predictor models showed that body size is negatively related to sexual dichromatism (Figure 6a, Table S13). Additionally, sexual dichromatism was more pronounced in more closed or forested habitats (Figure 6a, Table S13). In the multiple predictor models, the only effect that remained significant is that of body size on sexual dichromatism (Figure 6c, Table S15). The effect of habitat type on sexual dichromatism (Figure 6c, Table S15) was somewhat smaller and no longer significant, possibly due to reduced statistical power related to lower sample size (from N = 357 to N = 273). We found no effect of any of the predictors on the sexual difference in colour elaboration (Figure 6b and d, Table S14 and S16).

**Figure 6.**
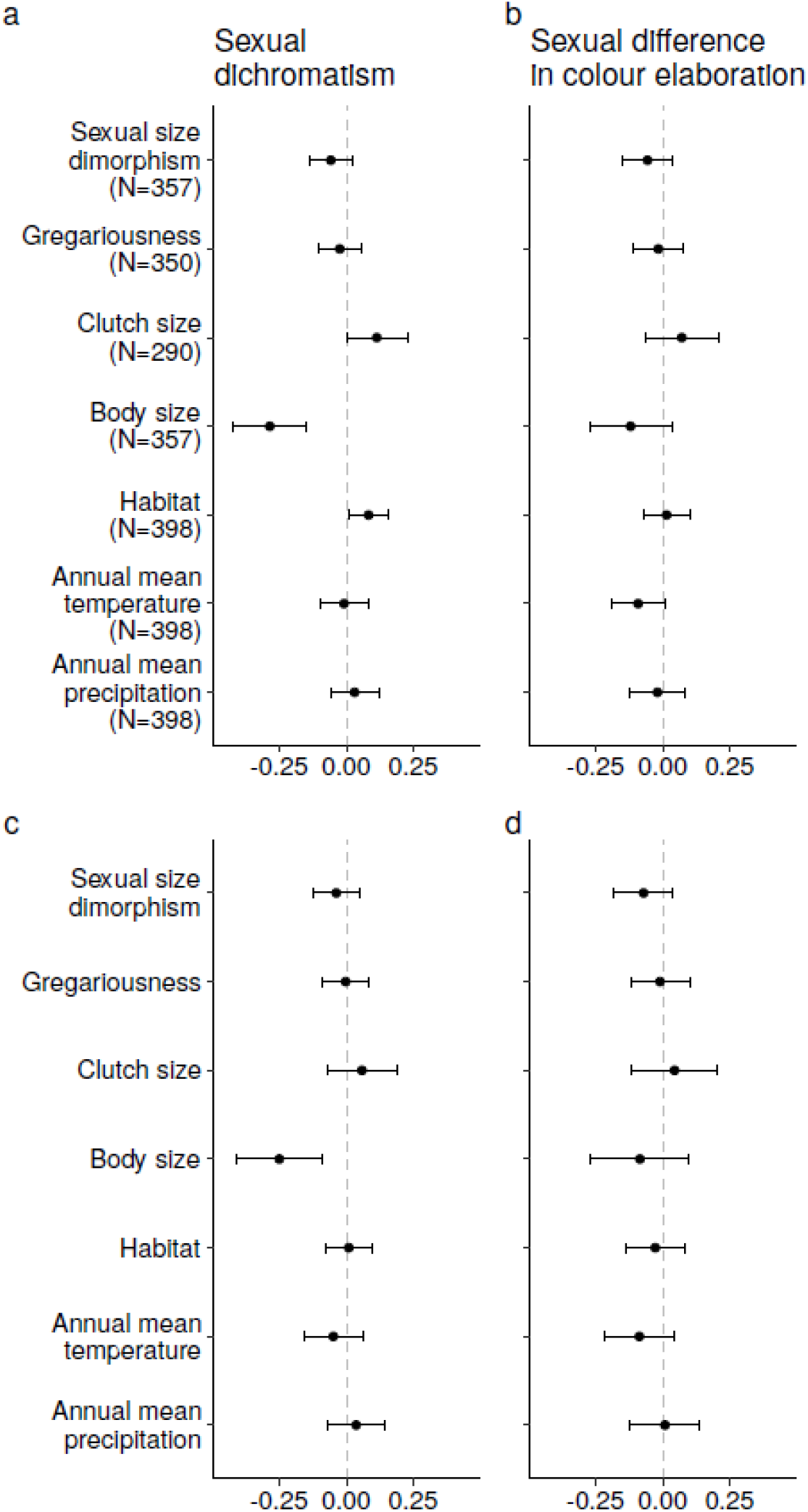
Effect sizes of predictors of difference in plumage colour between the sexes. Effect size of **a** sexual dichromatism and **b** sexual difference in colour elaboration based on single predictor models. Effect size of **c** sexual dichromatism and **d** sexual difference in colour elaboration based on multiple predictor models (N = 273 species). Shown are the means of the model coefficients for the 1000 phylogenetic linear models and the corresponding 95% confidence intervals. N indicates the number of species included in the analyses (determined by data availability).

### Variance explained by phylogeny

In all models, R^2^_*full*_ (variance explained by both phylogeny and fixed effects) was much higher (*range*: 0.274 – 0.669) than R^2^_*fixef*_ (variance explained only by the fixed effects, *range*: −1.57×10^-4^ – 0.21). This indicates that the phylogenetic signal in the residuals explains most of the variance in the models (see Table S1-S16).

## Discussion

Our study shows that variation in plumage colouration across all species of parrots, whilst strongly phylogenetically conserved, can be partly explained by key life-history traits and environmental variables. Among the former, body size seems the most important: larger species display more elaborate colours, such as red or blue, whilst smaller species had less elaborate plumage yet higher levels of sexual dichromatism (Figure 7 and Figure S3). Environmental effects were largely restricted to climatic variables and were partially in agreement with ecogeographical rules of colour variation. Two climatic variables correlate with plumage colour variation in parrots: temperature and precipitation.

**Figure 7.**
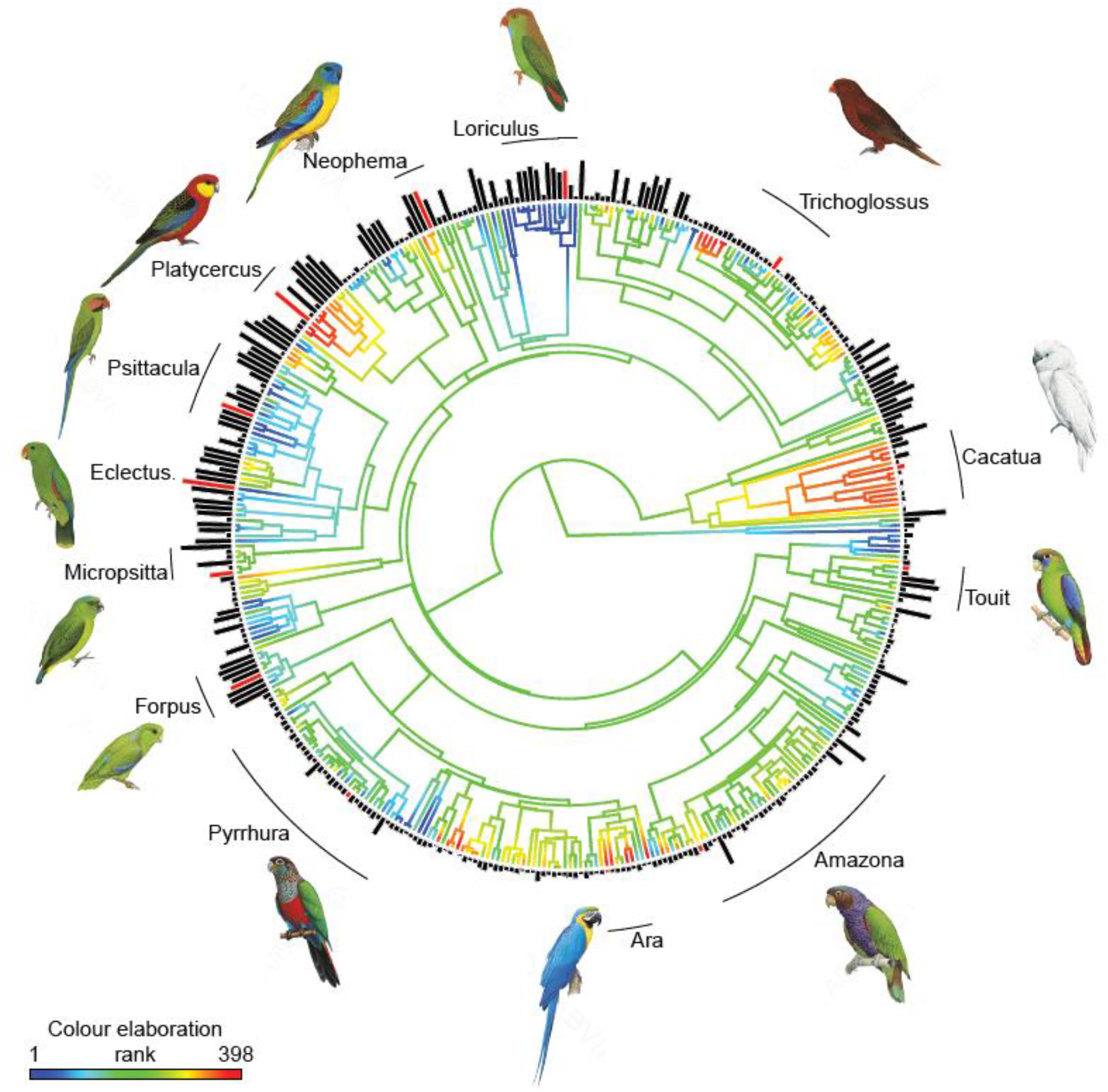
Parrots and cockatoos with more elaborate colours have lower levels of sexual dichromatism. Phylogeny of Psittaciformes depicting a reconstruction of evolutionary changes in male colour elaboration (branch colours, red = high, blue = low) using function *contMap* in R package “phytools” v.0.6-99 (Revell, 2012) and levels of sexual dichromatism (bar lengths at the tips). Note how species with low levels of colour elaboration have higher levels of sexual dichromatism. The plot is based on one phylogeny in the sample, but comparative analyses were carried out on 1000 phylogenetic reconstructions to account for phylogenetic uncertainty. Selected genera have been highlighted and species in illustrations are represented with red bars. Illustrations © Lynx Edicions.

Darker parrots are more frequent in humid environments, as predicted by Gloger’s rule (Rensch, 1936). Support for Gloger’s rule has already been found at the intraspecific level in parrots (in the crimson rosella *Platycercus elegans;* Ribot *et al*., 2019). We now show that it is a general pattern that applies at the interspecific level based on all 398 extant parrot species. There are two plausibly explanations for the correlation between humidity and darker colours (Delhey, 2017). First, darker colours would be favoured for camouflage in more humid environments as these harbour more vegetation and low light conditions. Second, as the presence of feather-degrading bacteria is higher in more humid environments, darker animals (with higher melanin concentration in their feathers) would be more resistant to feather degradation. Melanin deposition thickens the cortex of the barb and this makes feathers more resistant to feather-degrading bacteria (Bonser, 1995), which is more important in humid and warmer environments (Burtt & Ichida, 1999, 2004b).

Our results also show that males and females have more elaborated colours in warmer environments. As variation in temperature closely follows variation in latitude, this means that tropical parrots tend to be more colourful. Whether tropical birds are more colourful than their temperate counterparts has been a contested issue for nearly 200 years. Gloger, for example, suggested that tropical birds should be more pigmented and colourful because the environment was more benign allowing the production of such colours (Gloger, 1833). Proper tests of latitudinal patterns of colouration in birds have yielded conflicting results, some studies reporting no such correlation or even the opposite pattern (Bailey, 1978; Dalrymple *et al*., 2015), and others confirming the more elaborate colours of tropical species (Dale *et al*., 2015; Willson & von Neumann, 1972). Our findings agree with the latter, and are consistent with two non-mutually exclusive hypotheses (Dale *et al*., 2015). First, that tropical species are more colourful because mutual mate choice is stronger in those species; and second, because resource competition is stronger in the tropics, colour ornamentation might signal status in aggressive contexts. These effects are thought to be mediated by selection pressures associated with slow life histories typical of large species living in tropical environments.

We found that larger species display on average more elaborated colours, and also show darker, redder and more blue colours in their plumage. A similar finding has been reported in a large-scale comparative analysis of passerine plumage colour (Dale *et al*., 2015). Together, our results and those in Dale *et al*. (2015) disagree with the hypothesis that body size represents an evolutionary constraint on plumage colouration, as suggested by Galván *et al*. (2013). Firstly, Galván *et al*. (2013) suggested that larger species might be less colourful compared to smaller species because, proportionally to their size, the latter consume higher quantities of food (Tella *et al*., 2004). Hence, smaller species would have higher concentrations of limiting carotenoids pigments in their blood to colour their feathers. This explanation does not apply to parrots, since they do not deposit carotenoids in their plumage (Berg & Bennett, 2010). Secondly, they suggested that larger species might be able to detect other individuals at longer distances, whereas smaller species might have been forced to develop more conspicuous signals to communicate with conspecifics. Our results, on the contrary, are more consistent with the hypothesis that larger species experience lower predation pressure (Ricklefs, 2010), hence reducing selection for crypsis.

Our analyses further indicate that smaller parrot species –while displaying on average less elaborate colours– are more sexually dichromatic, in most cases (but not all) due to males having more elaborated colours than females (Figure S3). This suggests that smaller parrots are not only constrained from having highly elaborate colours, but also that the cost-benefit ratio of ornamental plumage colours varies between the sexes. Smaller species tend to have shorter lifespans (Bennett & Owens, 2002; de Magalhaes *et al*., 2007; Wasser & Sherman, 2010), which reduces the probability that a pair breeds together in subsequent seasons (Mauck, Marschall, & Parker, 1999). Under this scenario, higher levels of extra-pair paternity may be tolerated, i.e. it might not lead to reduced male investment, because males might invest more in current rather than in uncertain future reproduction (Mauck *et al*., 1999; Arnold & Owens, 2002). Previous studies showed that the frequency of extra-pair paternity is related to sexual dichromatism in birds (Møller & Birkhead, 1994; Owens & Hartley, 1998) and that dichromatic parrot species have longer sperm, and hence potentially higher levels of extra-pair paternity (Carballo *et al*., 2019). Thus, our finding that smaller parrot species are more dichromatic (with a tendency of males having more elaborated colours than females, Figure S3) may be a consequence of sexual selection via female choice for (extra-pair) mates. Sexual selection could also explain the observed relationship between habitat type and sexual dichromatism. Species inhabiting more forested habitats are more dichromatic possibly because bright colours would be favoured to help maximising conspicuousness of the sex under stronger sexual selection (Marchetti, 1993).

Many parrots form long-lasting pair bonds (Toft & Wright, 2015). Thus, larger species with longer lifespans (de Magalhaes *et al*., 2007; Wasser & Sherman, 2010) might be less dichromatic but display more elaborated colours as a consequence of mutual mate choice. As parrots are generally long-lived, especially compared with other bird species (Wasser & Sherman, 2010), we expect that both sexes are typically equally ornamented due to mutual mate choice, as observed in other tropical species (Bailey, 1978; Dale *et al*., 2015). The greater level of ornamentation parrots display (Delhey, 2015) might be due to mutual mate choice or the lack of selection on cryptic plumage in females that nest in cavities, at least in larger species. Moreover, the fact that suitable cavities are often a scarce resource may lead to strong competition between females (Heinsohn, Legge and Endler, 2005) for access to these resources and elaborate colouration may be selected as a signal of competitive ability or to advertise territory ownership.

In conclusion, our results are consistent with the idea that life-history traits reflecting predation pressure, the abiotic environment and possibly sexual selection have all shaped the evolution of plumage colouration in parrots. Body size had a consistent effect, indicating that this life-history trait plays a key role in the variation of colour elaboration and sexual dichromatism in parrots. Phylogenetic analyses indicated that an important component of the variation in parrot colouration and in sexual dichromatism was established in ancient evolutionary history, supporting results from comparative analyses in other birds (Brouwer & Griffith, 2019; Griffith, Owens, & Thuman, 2002). However, even though phylogeny explained most of the variation, we still found significant effects of life-history and environment on plumage colouration and sexual differences in parrots. Our comparative study leads to several testable hypotheses. First, we propose that larger species are more ornamented because of reduced selection against displaying colourful plumage given lower predation risk. Second, our results suggest that smaller species might experience more intense sexual selection on males, possibly via extra-pair paternity, whilst mutual mate choice might be common in larger species.

## Data Accessibility

All data, scripts and supplementary information accompanies this paper at https://osf.io/2xr4v/

## Acknowledgments

We thank Loro Parque, Loro Parque Fundación (LPF) and their respective presidents, Wolfgang and Christoph Kiessling, for the collaboration, the staff of LPF for support, Auguste M. P. von Bayern for developing the collaboration between the LPF and the Max Planck Institute for Ornithology, Pau Puigcerver and Rafael Zamora for support during data collection, and Laurie O’Neill for comments on manuscript drafts. Illustrations in figures were reproduced by permission of Lynx Edicions.

## Author Contributions

Conceived the study: L.C., M.V. and B.K. Collected the data: L.C. Analysed the data: L.C., M.V. and K.D. with input from B.K. Wrote the paper: L.C. with help of B.K and K.D. and input from M.V. L.C. is a member of the International Max Planck Research School (IMPRS) for Organismal Biology. This work was funded by the Max Planck Society (to B.K.).

