## Supplementary information for "Body size and climate as predictors of plumage colouration and sexual dichromatism in parrots"

### METHODS

#### Plumage score validation

This dataset contains reflectance measurements of 17 plumage patches (see Figure S1 in Delhey, 2015) from museum specimens of males (Nspecies = 50) and females (Nspecies = 49). We restricted measurements to those plumage patches that matched locations measured from bookplates (this excludes upper and lower back, rump, cheek, belly and vent). Reflectance spectra, encompassing the entire visual range of birds (300-700 nm), were summarised using psychophysical models of avian vision (Vorobyev & Osorio, 1998) using formulas in Cassey et al. (2008) as implemented in R by Delhey, Delhey, Kempenaers, & Peters (2015). These models require knowledge of the visual sensitivity of the photoreceptors in the retina, the four single cones that mediate colour vision (VS, S, M and L, sensitive to very short, short, medium and long wavelengths, respectively) and the double cones which are involved in perception of achromatic (i.e. light to dark) variation (Cuthill, 2006). Whilst the sensitivity function of double cones does not seem to vary much between species, the sensitivity function of VS and S single cones does. Species thus fall into two main groups: U-type and V-type species, whereby functions for U-type species peak deeper into the UV range than those of V-type species (even though the latter have some UV sensitivity, Hart & Hunt, 2007). Psittaciformes seem to all have U-type visual sensitivities (Ödeen & Håstad, 2013)and we used these as obtained from Endler & Mielke (2005). As illuminant we used the spectrum of standard daylight (d65). We estimated the noise-to-signal ratio of each photoreceptor by using the average abundance of each single cone type in the retina as obtained from Hart (2001), a Weber fraction of 0.1 for the L cone (Olsson, Lind, & Kelber, 2018) and formula 10 in Vorobyev, Osorio, Bennett, Marshall, & Cuthill (1998). For the double cones the noise-to-signal ratio used was 0.2 (Olsson et al., 2018).

Using this approach visual models yield, for each reflectance spectrum, three chromatic variables xyz and one achromatic variable L (light to dark variation), note that there is one more variable than in the CIELAB space because birds have four types of cones involved in colour vision and humans only three. We can then use the output of visual models to compute colour elaboration, sexual dichromatism, differences in colour elaboration between males and females and colour diversity as described above from bookplates.

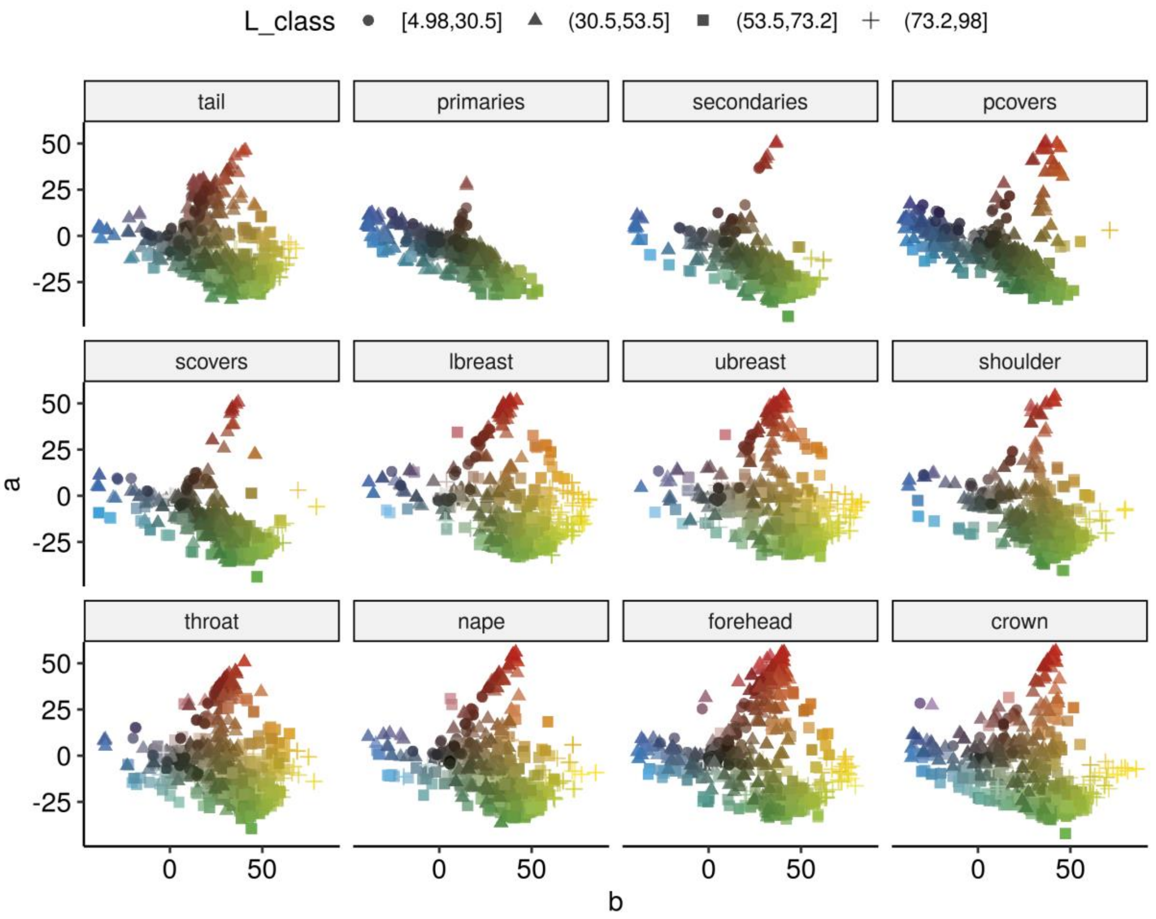


**Figure S1**. Colour distribution of the raw data extracted from 398 parrot species by body patches. Colour space was measured in the three CIELAB coordinates (*L*,*a* and *b*).

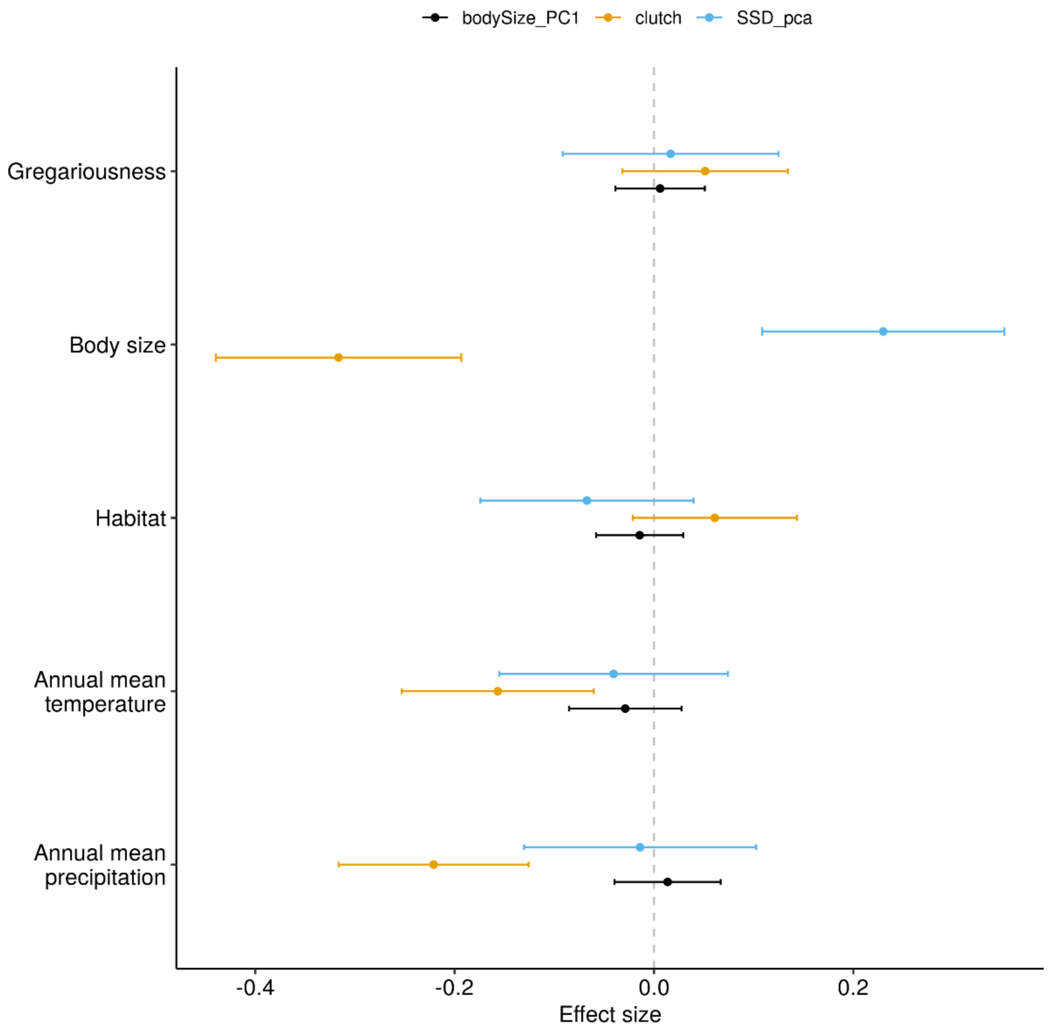


**Figure S2**. Effect sizes for each of the predictor variables on sexual size dimorphism (SSD), clutch size and body size based on multiple predictor models (N = 272 species for clutch size and N = 324 for SSD and body size, Table S17-S19). Black denotes body size, orange clutch size and blue SSD. Shown are the means of the model coefficients for the 1000 phylogenetic linear models and the corresponding 95% confidence intervals.

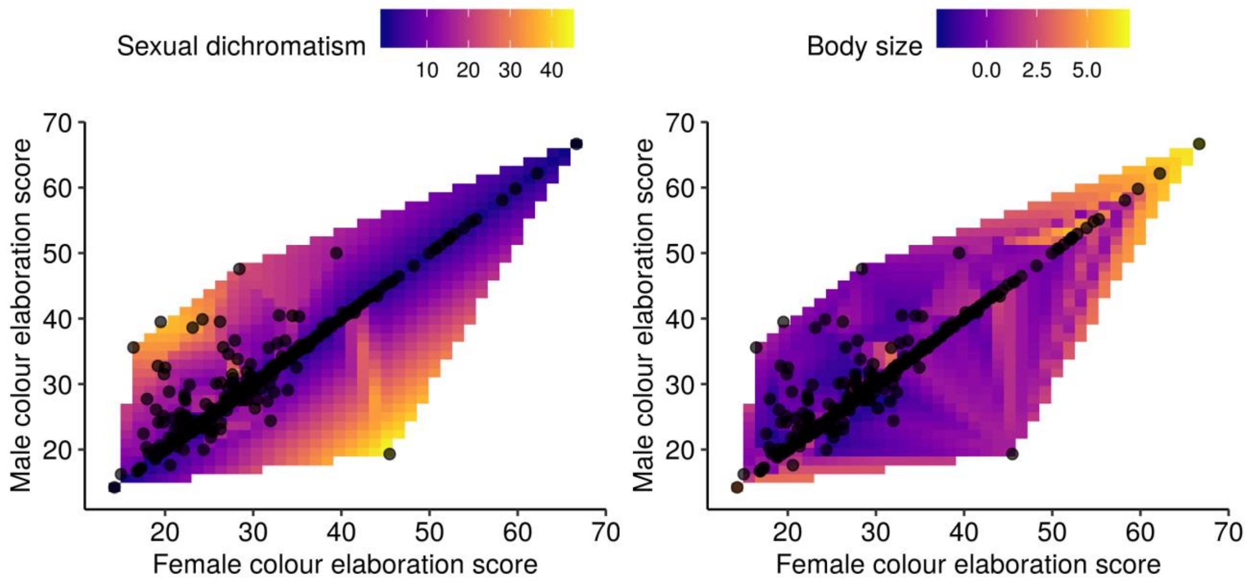


**Figure S3**. Male colour elaboration score versus female colour elaboration score. Heatmaps show variation of the sexual dichromatism score (left panel) and body size (right panel). Black dots represent distribution of the raw data extracted from 398 parrot species.
